## Supplementary Information for "Uptake of small extracellular vesicles by recipient cells is facilitated by paracrine adhesion signaling"

This PDF file includes:

Supplementary Figs. 1 to 19

Captions for Supplementary movies 1 to 7

Supplementary Table 1

Supplementary References

**Supplementary Figures**

Supplementary Figure 1

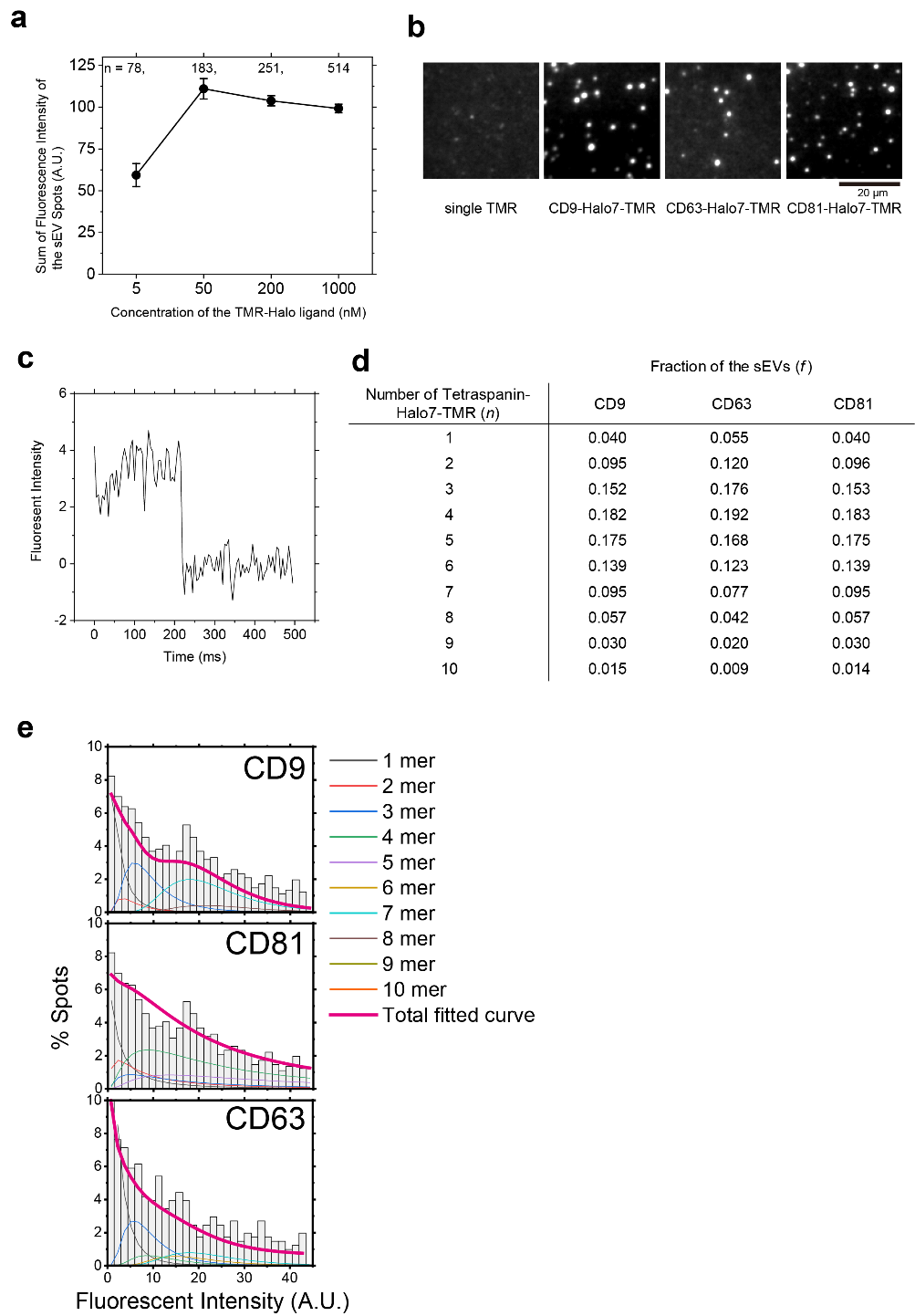

**Supplementary Fig. 1.** Tetraspanin marker proteins in sEVs were labeled with TMR via Halo7 at high efficiency. **a**, Sum of the fluorescence intensities of the individual spots of sEVs containing CD63-Halo7 labeled with various concentrations of TMR Halo ligands after incubation for 1 h. The fluorescently labeled tetraspanin marker proteins in sEVs were observed by TIRFM. The elevation of TMR-Halo ligand concentration above 50 nM did not increase the sum of the fluorescence intensities. As shown in the previous reports^1, 2^, these results indicate that the tetraspanin marker proteins were labeled with TMR via Halo7 at high efficiency. “n” indicates the number of examined particles. **b**, Typical images of single molecules of TMRs and individual PC-3 cell-derived sEV particles containing CD9-Halo7-TMR, CD63-Halo7-TMR, and CD81-Halo7-TMR. The expression levels of the tetraspanin-Halo7 proteins in sEVs derived from PC-3 cells are shown in Fig. 1c. **c**, A single-step photobleaching event of a TMR molecule (shown in Supplementary Fig. 1b) at 215 ms. We have previously validated the linear relationship between the number of molecules in clusters and fluorescence intensity in our previous studies^1-3^. **d**, Table shows the fractions (*f*) of sEVs containing the numbers (*n*) of tetraspanin-Halo7-TMR molecules indicated in the left column, which were estimated by fitting the histograms of fluorescence intensities of TMR in sEV particles with the lognormal function curves. The average numbers of tetraspanin-Halo7-TMR molecules per sEV particle were calculated by summing the values of *f* × *n*. **e**, Distribution of TMR fluorescence intensities from TSN-Halo7-labeled individual sEV particles. The data were fitted with ten log-normal functions (shown in different colors).

Supplementary Figure 2

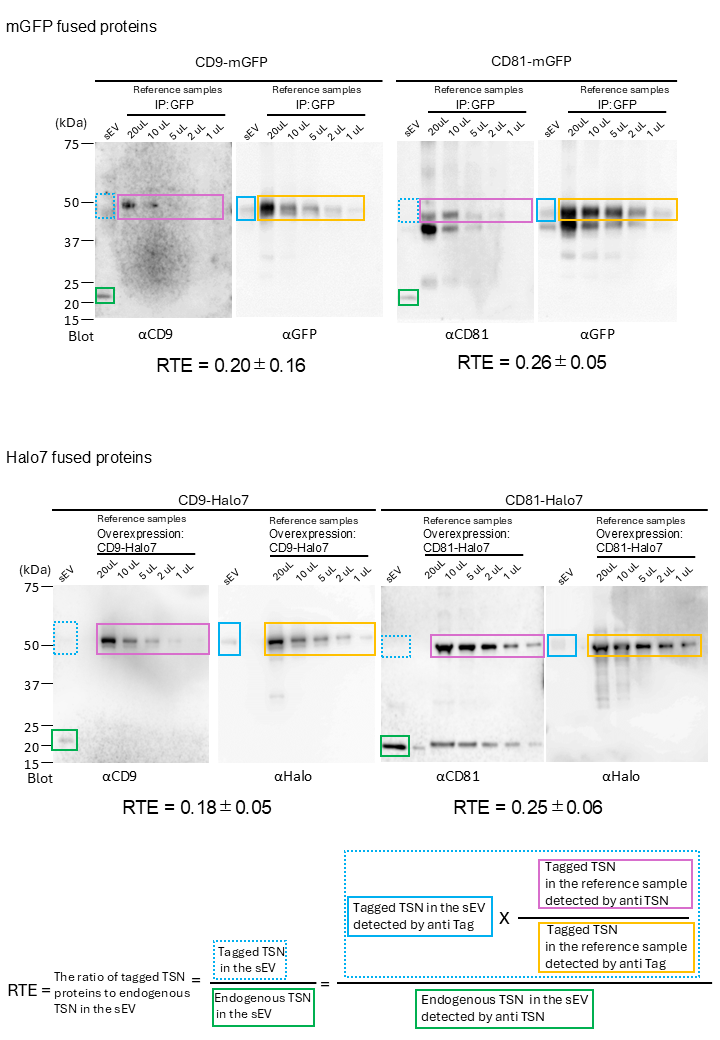

**Supplementary Fig. 2.** Estimation of the ratio of tagged tetraspanins (TSN), CD9 and CD81 to their endogenous counterparts in sEVs (RTE). To compare the quantities of tagged TSN proteins with their endogenous counterparts, we prepared reference samples for band intensity calibration. mGFP- or Halo7-tagged CD9 or CD81 was concentrated to detectable levels by immunoprecipitation (upper) or overexpression (bottom). Note that these concentrated reference samples serve exclusively as intensity references for band detection, rather than as indicators of protein sorting efficiency into EVs. First, using the reference samples, we calculated the ratio of band intensities of tagged TSN proteins detected with anti-TSN antibodies (purple box) to those detected with anti-tag antibodies (yellow box), which remains independent of protein sorting efficiency into EVs. Next, we quantified the band intensity of tagged TSN proteins in sEVs using anti-tag antibodies (light-blue box). This value was then multiplied by the previously calculated ratio to estimate the theoretical band intensity of marker proteins below the detection limit (dotted light-blue box). Finally, this calculated value was divided by the band intensity of endogenous proteins in sEVs detected with anti-TSN antibodies (green box) to determine the ratio of tagged marker proteins to endogenous ones (RTE), as shown in the formula below. The RTE values for CD9 and CD81 ranged from 0.18 to 0.26, indicating that these tagged proteins were expressed at much lower levels than their endogenous counterparts. All blots were quantified from three independent experiments.

Supplementary Figure 3

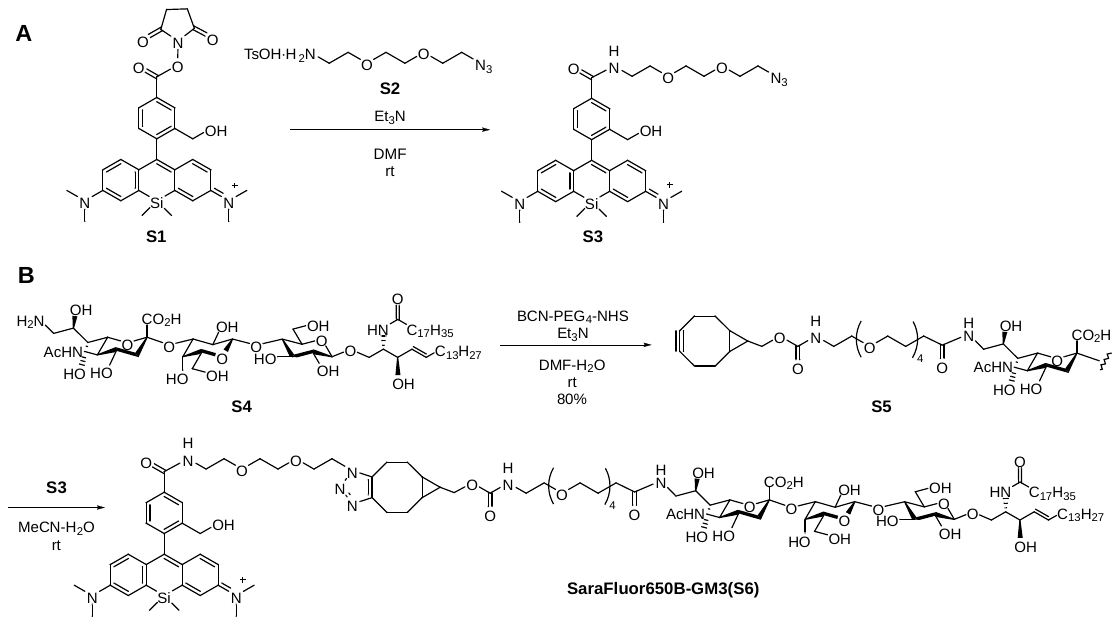

**Supplementary Fig. 3. Synthesis of SaraFluor650B-GM3**

**a,** Synthesis of S5. S4 (3.2 mg, 2.7 μmol) and endo-BCN-PEG_4_ *N*-succinimidyl ester (BroadPharm, (California, USA)) (2.2 mg, 4.1 μmol) were dissolved in DMF/H_2_O (499 µL/45 µL) at room temperature. Et_3_N (7.6 µL, 54 μmol) was added at room temperature. The reaction mixture was stirred for 1 h at room temperature as the reaction was monitored by TLC (CHCl_3_/MeOH/5% CaCl_2_ aq. = 5:3:0.25). The reaction mixture was directly subjected to column chromatography on Sephadex LH-20 using CHCl_3_/MeOH (1:1) as the eluent and column chromatography on Iatrobeads using (CHCl_3_/MeOH/H_2_O = 2:1:0 to 1:1:0 to 5:4:1) to give BCN-GM3 (S5) (3.5 mg, 80%): [α]_D_ −19.0 (c 0.3, MeOH); ^1^H NMR (800 MHz, CD_3_OD) δ 5.68 (m, 1 H, *J*_5,6a_ = 6.4 Hz, *J*_5,6b_ = 8.0 Hz, *J*_4,5_ = 14.4 Hz, H-5*^Cer^*), 5.44 (dd, 1 H, *J*_3,4_ = 7.2 Hz, H-4*^Cer^*), 4.42 (d, 1 H, *J*_1,2_ = 8.0 Hz, H-1), 4.30 (d, 1 H, *J*_1,2_ = 8.0 Hz, H-1), 4.19 (dd, 1 H, *J*_1a,2_ = 4.0 Hz, *J*_gem_ = 9.6 Hz, H-1a*^Cer^*), 4.14 (d, 2 H, NHCOOC*H*_2_), 4.07 (t, 1 H, *J*_2,3_ = 8.0 Hz, H-3*^Cer^*), 4.03–3.25 (m, 39 H, H-4*^Neu^*, H-5*^Neu^*, H-6*^Neu^*, H-7*^Neu^*, H-8*^Neu^*, H-9a*^Neu^*, H-9b*^Neu^*, H-2*^Gal^*, H-3*^Gal^*, H-4*^Gal^*, H-5*^Gal^*, H-6a*^Gal^*, H-6b*^Gal^*, H-2*^Glc^*, H-3*^Glc^*, H-4*^Glc^*, H-5*^Glc^*, H-6a*^Glc^*, H-6b*^Glc^*, H-1b*^Cer^*, H-2*^Cer^*, 9 CH_2_), 2.85 (m, 1 H, H-3*eq^Neu^*), 2.53–2.47 (m, 2 H, NHCOC*H*_2_), 2.27–2.16 (m, 8 H, NHCOC*H*_2_*^Cer^*, 6 H*^BCN^*), 2.04–2.01 (m, 5 H, H-6a*^Cer^*, H-6b*^Cer^*, Ac), 1.73 (m, 1 H, H-3*ax^Neu^*), 1.61–1.29 (m, 55 H, 26 CH_2_*^Cer^*, 3 H*^BCN^*), 0.96–0.89 (m, 8 H, 2 CH_3_*^Cer^*, 2 H*^BCN^*); ^13^C NMR (200 MHz, CD_3_OD) δ 175.9, 175.4, 174.9, 174.2, 159.3, 135.0, 131.4, 105.1, 104.6, 101.2, 99.6, 80.9, 77.7, 77.1, 76.5, 76.3, 74.9, 74.8, 73.0, 72.0, 71.5, 71.5, 71.4, 71.3, 71.3, 71.1, 71.0, 70.9, 70.0, 69.4, 69.0, 68.4, 63.7, 62.8, 61.9, 54.7, 53.9, 43.9, 42.0, 41.7, 37.5, 37.4, 33.5, 33.1, 33.1, 30.9, 30.9, 30.8, 30.8, 30.8, 30.7, 30.7, 30.5, 30.5, 30.5, 30.5, 30.2, 27.2, 23.8, 22.6, 22.0, 21.4, 19.0, 14.5; HRMS (ESI) *m/z*: found [M−H]^−^ 1601.9789, C_81_H_142_N_4_O_27_ calcd for [M−H]^−^ 1601.9789.

**b,** Synthesis of S6 (Preparation of S3). S1 (Goryo Chemical, Inc., (Sapporo, Japan))(ca. 50 μg, ca. 0.090 μmol) and azido-PEG_2_-amine·Tos-OH S2 (0.1 mg, 0.3 μmol) were dissolved in DMF (36 µL) at room temperature. Et_3_N (1.0 µL, 7.2 μmol) was added at room temperature. After stirring for 2 h at room temperature, azido-PEG_2_-amine·Tos-OH S2 (0.1 mg, 0.3 μmol) was added. After stirring for 10 min at room temperature as the reaction was monitored by TLC (CHCl_3_/MeOH = 40:1), the reaction mixture was concentrated, and the residue was purified by column chromatography on Iatrobeads using CHCl_3_/MeOH (100:1 to 1:1) as the eluent to give SaraFluor650B-linkerN_3_ (S3): ^1^H NMR (800 MHz, CD_3_OD) δ 7.82 (s, 1 H, Ar), 7.64 (d, 1 H, Ar), 7.01–6.89 (m, 5 H, Ar), 6.71 (dd, 2 H, Ar), 5.40 (s, 2 H, C*H*_2_OH), 3.67–3.57 (m, 12 H, 6 CH_2_), 2.93 (s, 12 H, 4 NCH_3_), 0.59 (s, 3 H, SiCH_3_), 0.49 (s, 3 H, SiCH_3_); ^13^C NMR (200 MHz, CD_3_OD) δ 170.1, 163.7, 150.6, 139.8, 139.4, 135.5, 135.1, 130.0, 128.0, 124.8, 117.6, 115.7, 94.2, 71.5, 71.4, 71.1, 70.5, 51.7, 40.8, 30.8, 30.8, 0.2, −0.3; HRMS (ESI) *m/z*: found [M+Na]^+^ 637.2926, C_33_H_42_N_6_O_4_Si calcd for [M+Na]^+^ 637.2929.

(Click reaction)

A half amount of prepared S3 and S5 (0.2 mg, 0.1 μmol) were dissolved in MeCN/H_2_O (200 µL/100 µL) at room temperature. After stirring for 19 h at room temperature as the reaction was monitored by TLC (CHCl_3_/MeOH/5% CaCl_2_ aq. = 5:3:0.2), the reaction mixture was concentrated. The residue was purified by PTLC using (CHCl_3_/MeOH/H_2_O = 5:3:0.2) to give SaraFluor650B-GM3 (S6): HRMS (ESI) *m/z*: found [M−H]^−^ 2216.2729, C_114_H_184_N_10_O_31_Si calcd for [M−H]^−^ 2216.2825.

Supplementary Figure 4

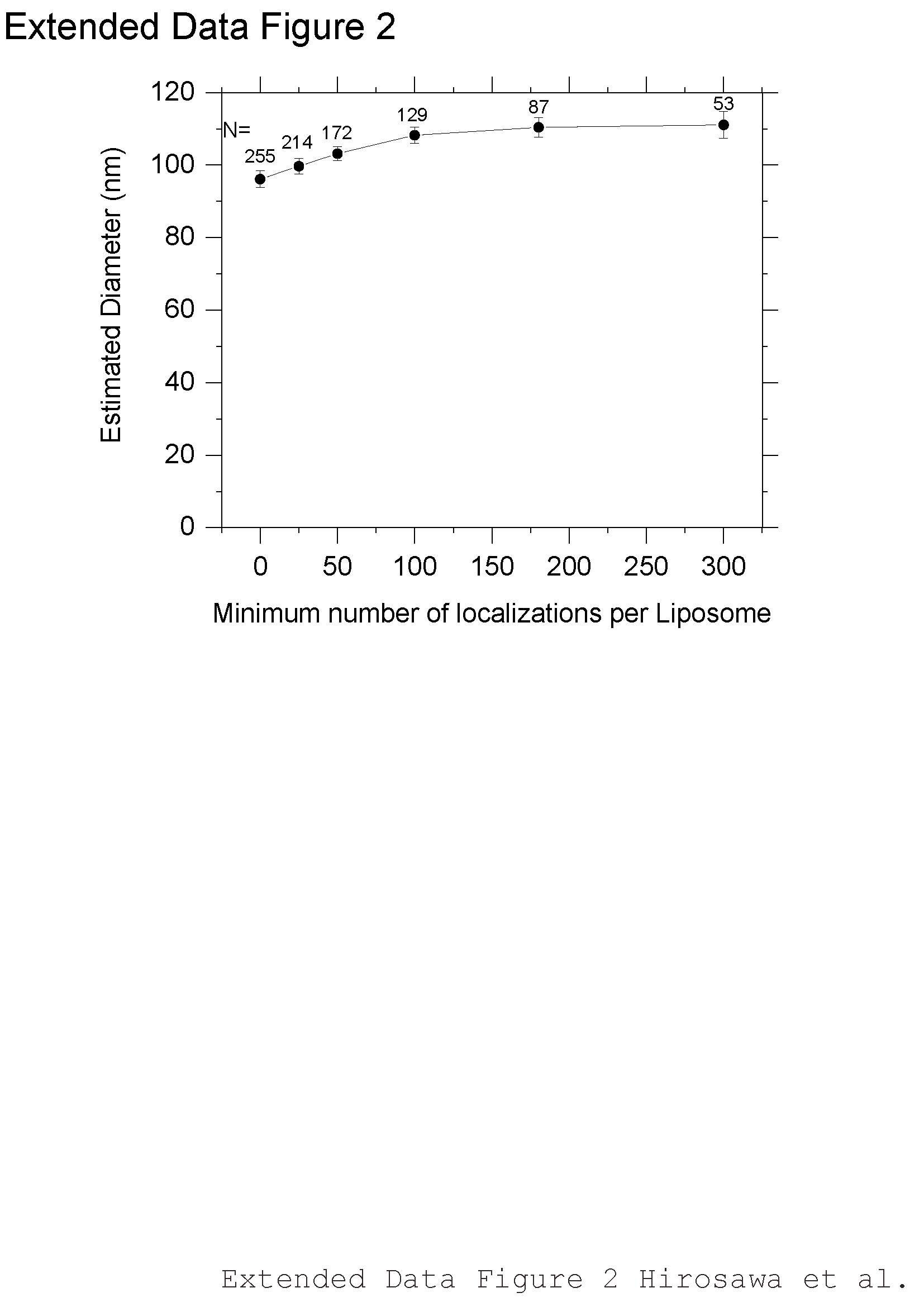

**Supplementary Fig. 4.** Estimation of the liposome diameter by dSTORM

The liposome diameter was estimated via the Voronoi method. Each number of localizations was obtained by dSTORM observation of SF650B-GM3 in liposomes. The x-axis indicates the minimum number of localizations used to estimate the diameter of each liposome. The estimated diameter was saturated above 100 localizations. “N” indicates the number of examined liposomes. The data are presented as the means ± SE.

Supplementary Figure 5

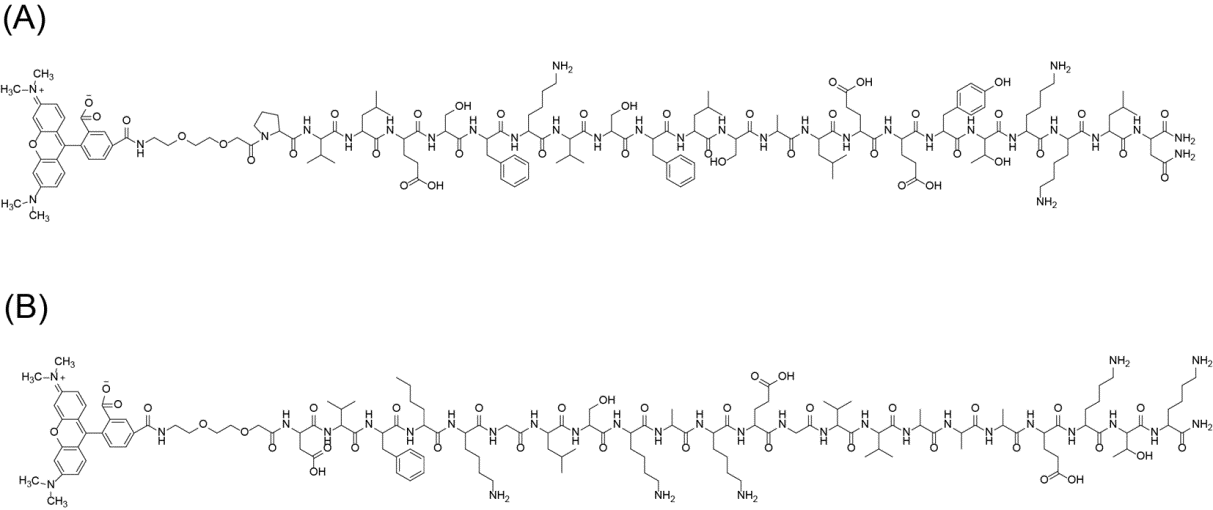
**a**

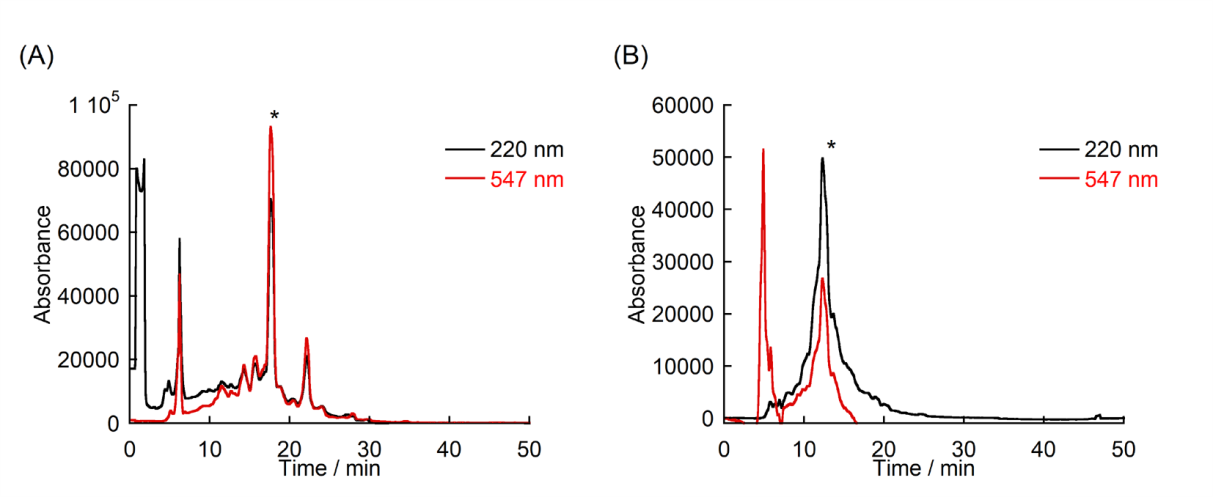
**b**

**c**

|  | Observed mass (m/z) | Calculated mass [M+H]^+^ | Retention time |
| --- | --- | --- | --- |
| ApoC-TAMRA | 3097.28 | 3097.61 | 17.5 min |
| α-synuclein(p2-23)-TAMRA | 2845.89 | 2845.54 | 13.0 min |

**Supplementary Fig. 5. a.** Chemical structure of ApoC-TAMRA **(A)** and α-synuclein(p2-23)-TAMRA. **(B)**. **b.** HPLC profiles for the purification of probes (ApoC-TAMRA **(A)** and α-synuclein(p2-23)-TAMRA **(B)**). Gradient condition: **(A)** 35-70% CH_3_CN (0.1% TFA) in H_2_O (0.1% TFA) during 50 min, **(B)** dd-60 CH_3_CN (0.1% TFA) in H_2_O (0.1% TFA) during 50 min. Absorbance at 220 nm (peptide) and 547 nm (TAMRA) was monitored. The peak (*) was for collected and identified as the purified probe by MALD-TOF-MS. **c.** Probe characterization

Supplementary Figure 6

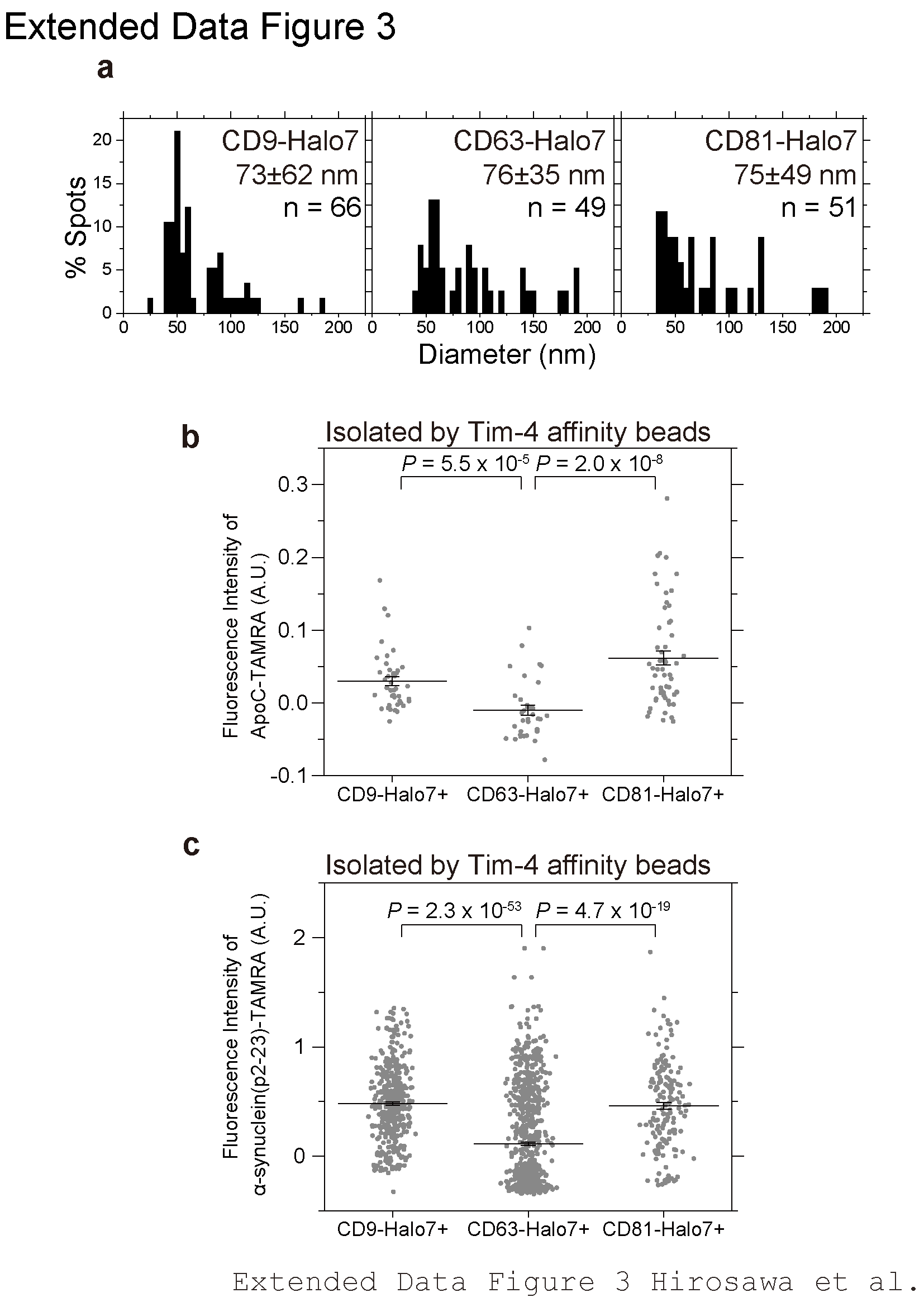

**Supplementary Fig. 6.** sEV subtypes isolated by the Tim-4 bead method exhibit different membrane defects. **a**, Diameter distributions of sEVs isolated by Tim-4 beads, which were determined by dSTORM methods. The data are presented as the means ± SD. **b**, **c**, Distributions of the fluorescence intensities of ApoC-TAMRA (**b**) and amphipathic helix peptide (p2-23) of α-synuclein (**c**) on the individual sEV particles that contained CD9, CD63, and CD81 labeled with SF650T via Halo7; these particles were isolated by the Tim-4 bead method and observed by TIRFM. “n” indicates the number of examined particles. The data are presented as the means ± SE, and the statistical tests were performed with Welch’s *t*-test (two-sided).

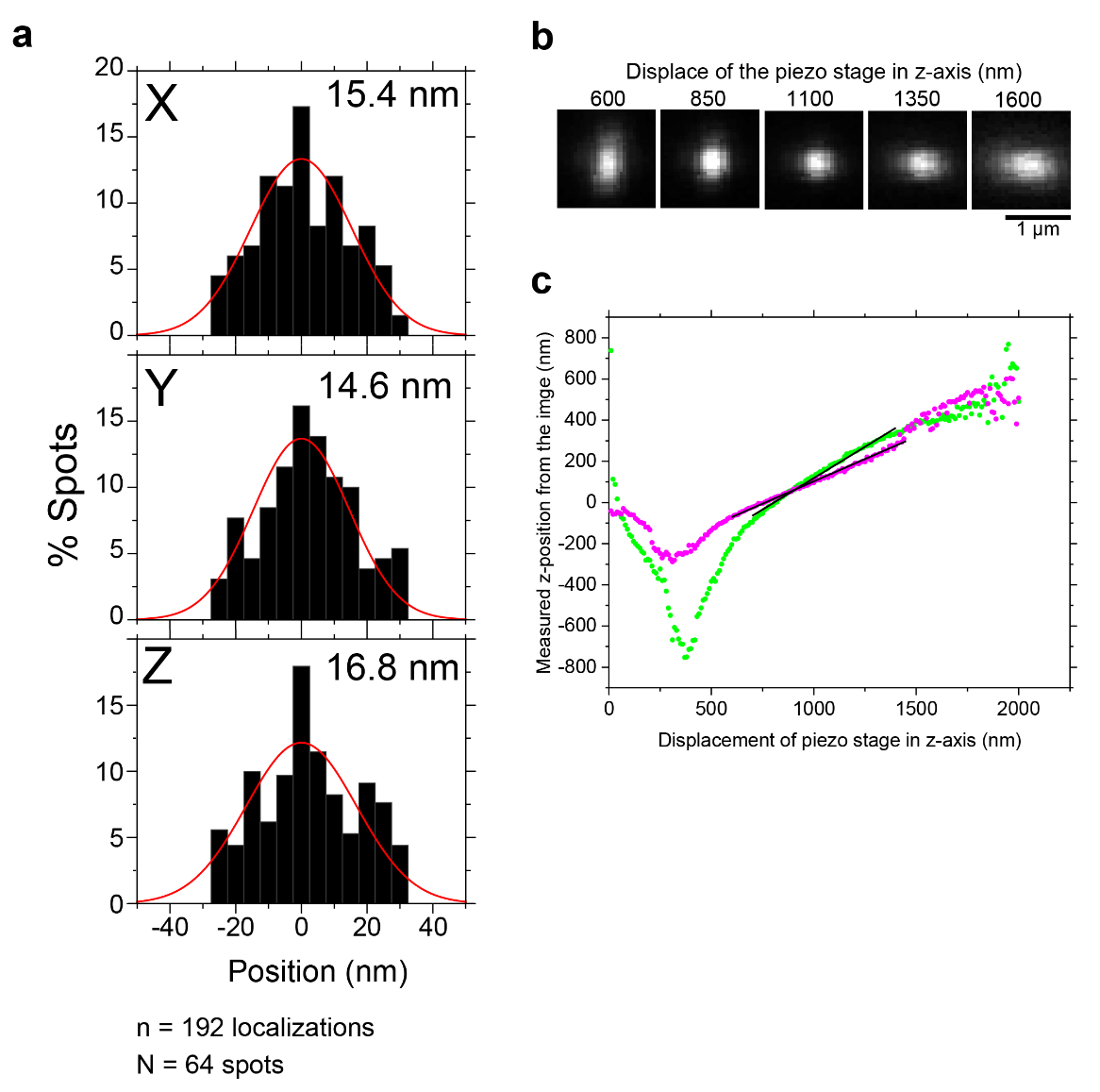
Supplementary Figure 7

**Supplementary Fig. 7.** **a**, Accuracy of the position determined for single sEV particles

sEV-CD63Halo7-SF650T particles were attached to a coverslip, observed by oblique angle illumination, and tracked in 3D. The center position and eccentricity were measured every 33 ms for 99 ms using ThunderSTORM software. Histograms of the displacements (n= 192 localizations) in the x-, y- and z-directions for all of the spots (N= 64) on the images were obtained. The histograms were fitted by Gaussian curves. The standard deviations in x-, y- and z-directions were 15.4, 14.6, and 16.8 nm, respectively. **b**, Validation of 3-D single-particle tracking. Astigmatic images of 100-nm-diameter multicolor fluorescent beads attached to a coverslip. Images were captured by oblique angle illumination at 250 nm intervals along the Z-axis during piezo stage elevation. **c**, Z-position linearity analysis. Fluorescent beads were imaged during piezo stage elevation (1 nm/ms). Z-positions were estimated from the image ellipticity as described in Methods^4^. Green and magenta symbols represent data from images of short wavelength (Ex. 560 nm/Em. 580 nm) and long wavelength (Ex. 660 nm/Em. 680 nm) fluorescent spots, respectively. Solid black lines show linear regression fits.

Supplementary Figure 8

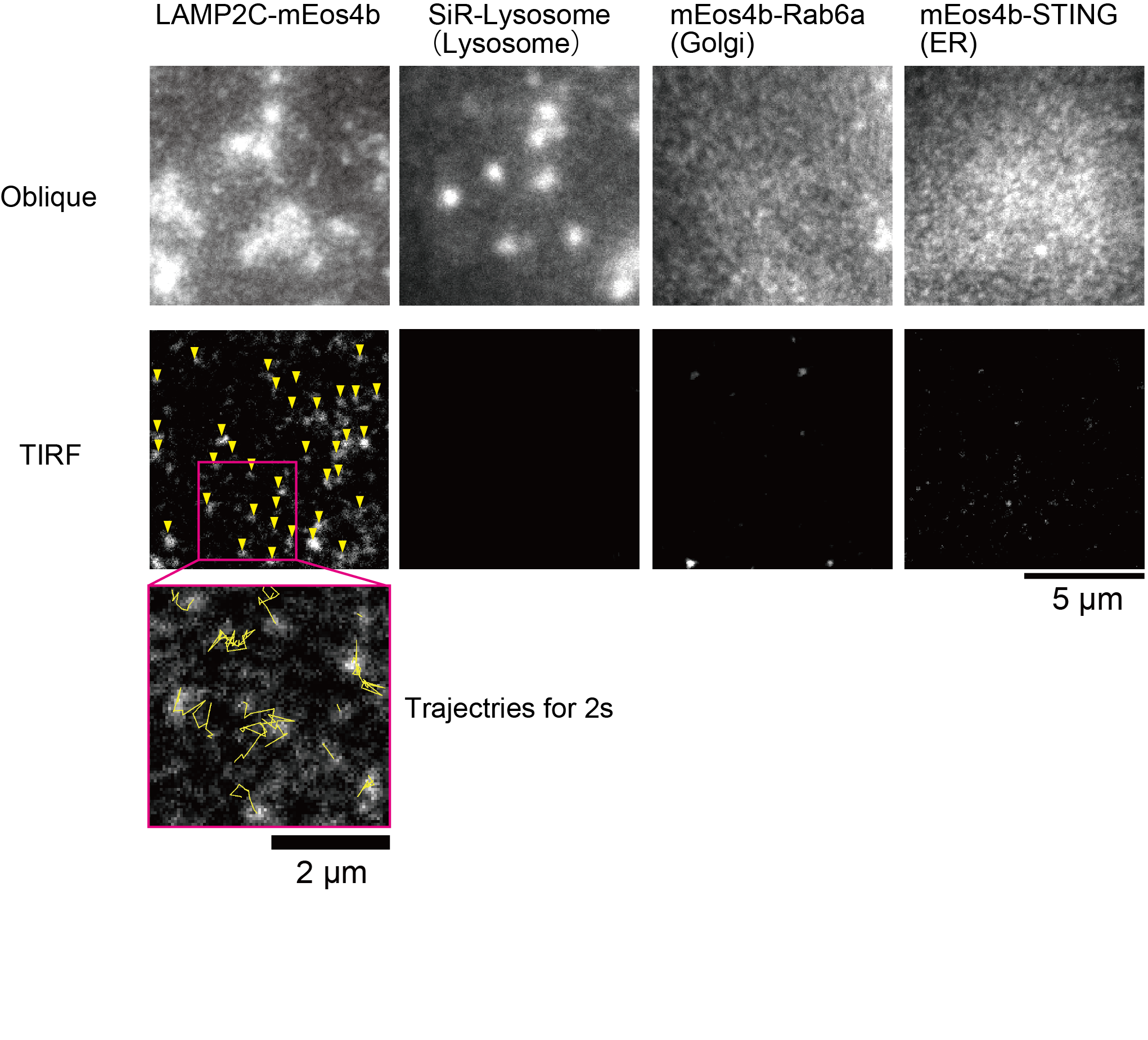

**Supplementary Fig. 8.** Fluorescent images of LAMP2C-mEos4b, SiR-Lysosome, mEos4b-STING, and mEos4b-Rab6a in PZ-HPV-7 cells. Fluorescent images were acquired by oblique angle illumination (top panels) and TIRFM (middle panels). The single molecules of LAMP2C -mEos4b are indicated by yellow arrowheads (middle, most left panel). Trajectories of single fluorescent molecules of LAMP2C -mEos4b observed by TIRFM are shown in the bottom panel. Scale bar: 10 μm.

Supplementary Figure 9

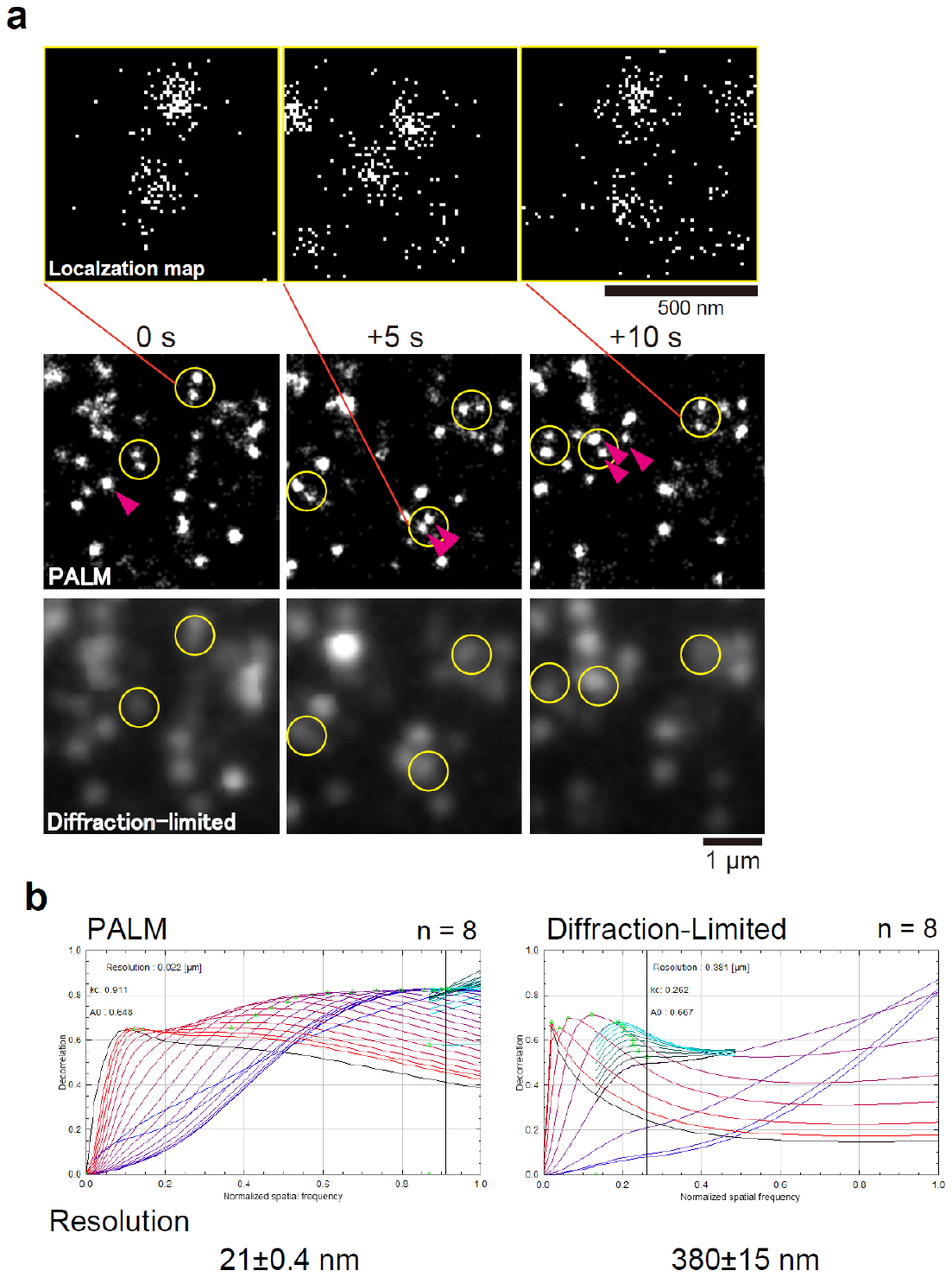

**Supplementary Fig. 9.** PALM image sequence obtained to track the dynamics of membrane structures. **a**, Typical PALM image sequence of caveolin-1 (Cav1)-mEos4b in PZ-HPV-7 cells. Single molecules of Cav1-mEos4b were observed by TIRFM at 5 ms/frame for 1002 frames (see Fig. 4b). The center coordinates of the spots were plotted on a localization map (top). Subsequently, the distributions of each localization, including their uncertainties, were combined to create a PALM image (middle). Caveolae, highlighted by yellow circles, can be resolved as two distinct structures in the PALM images, whereas they appear as single entities in the diffraction-limited images (bottom). Caveolae marked by magenta arrowheads did not exist at the same position in the previous and/or next frame. **b**, Parameter-free image decorrelation analyses with Ng=50 and Nr=50 ^5^. Vertical gray lines: locations of kc, which is the cutoff frequency. Ng is the number of high-pass filters used to determine the resolution, and Nr is the number of points used to calculate the normalized frequency plots. The average spatial resolutions of 8 PALM images and 8 diffraction-limited images of caveolae were estimated to be 21 ± 0.4 nm (mean ± SE) and 380 ± 15 nm, respectively.

Supplementary Figure 10

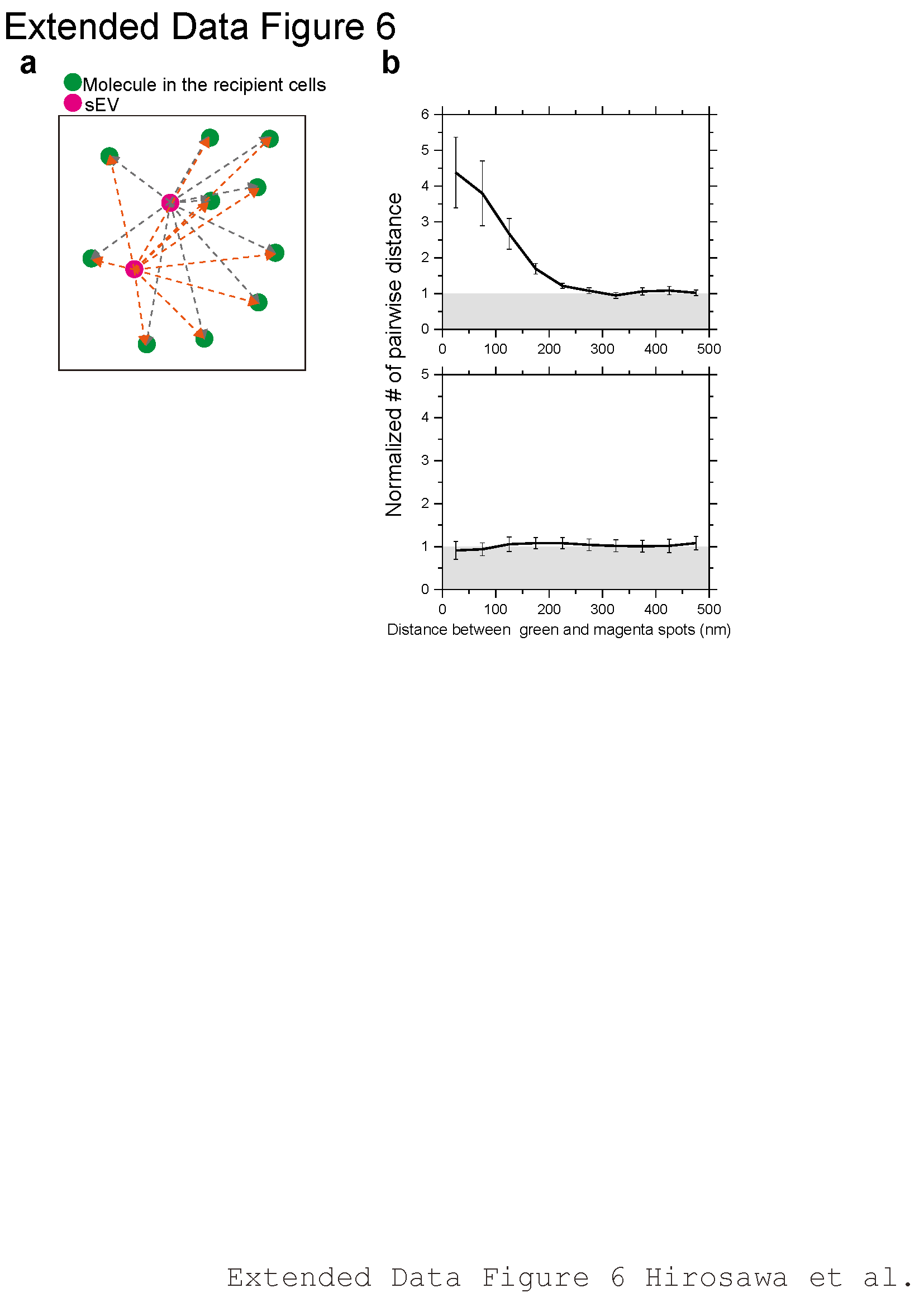

**Supplementary Fig. 10.** Correlation analysis of the fluorescent spots. **a**, In the movies showing simultaneous dual-color single-molecules, the spatial pair correlation function was determined and plotted in a histogram, showing the number densities of pairwise distances (the number of pairs normalized with the distance) at a given distance^6^. First, a region of interest (ROI) greater than 10 μm was selected, the distances between all the pairs of green and magenta spots in the ROI in a video frame were measured, and this process was repeated for all video frames in an image sequence. **b**, Distribution of the number of distances normalized for the unit area (i.e., the number density) for all the pairs was plotted against the distance. A bin size of 50 nm and an overall x-axis size of 500 nm were used throughout this study. This process was repeated for all sets of green and magenta videos, and the mean ± SE values were plotted (top). As a control, the magenta image was superimposed on the 180-degree rotated green image (bottom).

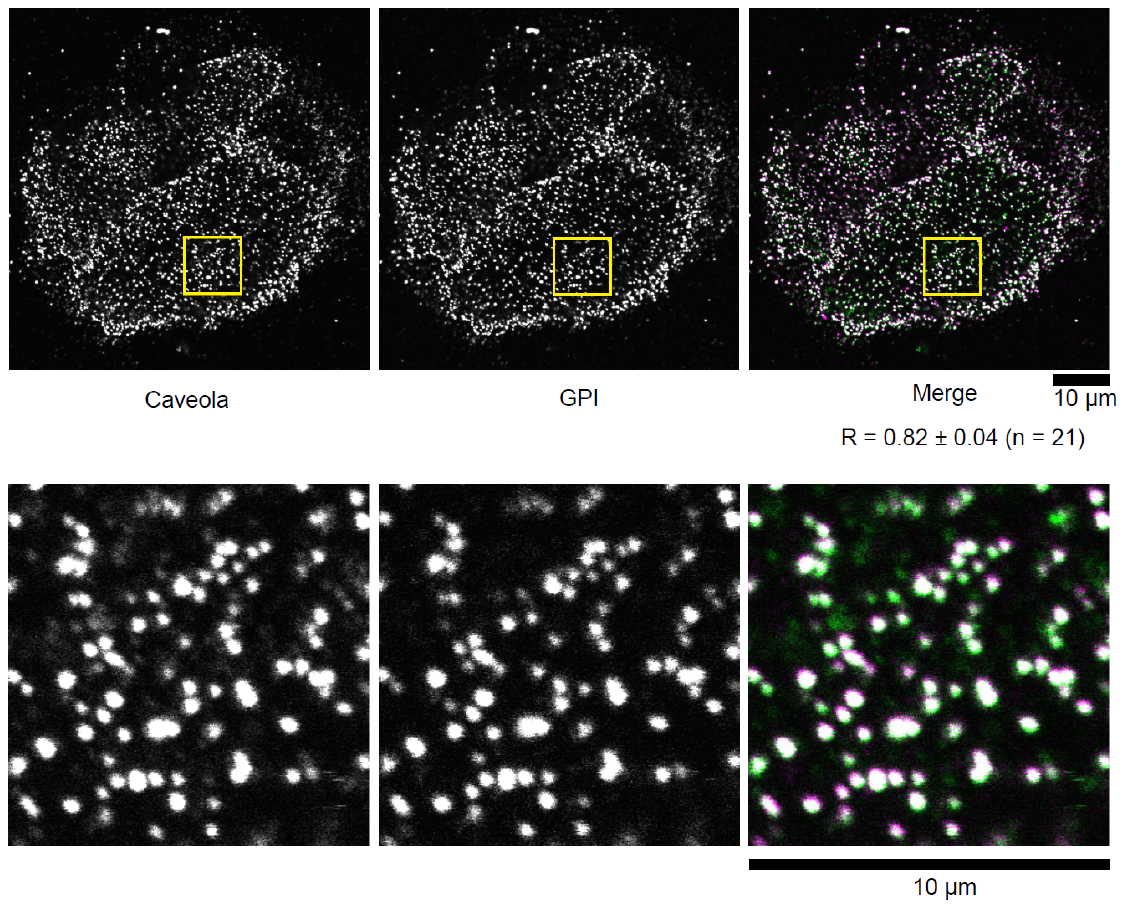
Supplementary Figure 11

**Supplementary Fig. 11.** Crosslinked GPI-anchored proteins were colocalized with caveolae. PZ-HPV-7 cells expressing Halo-GPI were incubated with rabbit polyclonal anti-Halo antibodies (10 μg/mL) for 10 min at 37°C, followed by incubation with rhodamine-labeled secondary antibody for rabbit IgG for 10 min at 37°C to crosslink Halo-GPI. The cells were then fixed with 4% paraformaldehyde for 90 min at room temperature. Caveolae were labeled using mouse anti-caveolin-1 antibody, followed by Alexa488-labeled secondary antibody for mouse IgG. Fluorescence images were acquired by confocal microscopy. Pearson's correlation coefficient was 0.82 ± 0.04 (mean ± SE, n = 21 cells). The magnified images of the yellow square in the top panels are shown at the bottom.

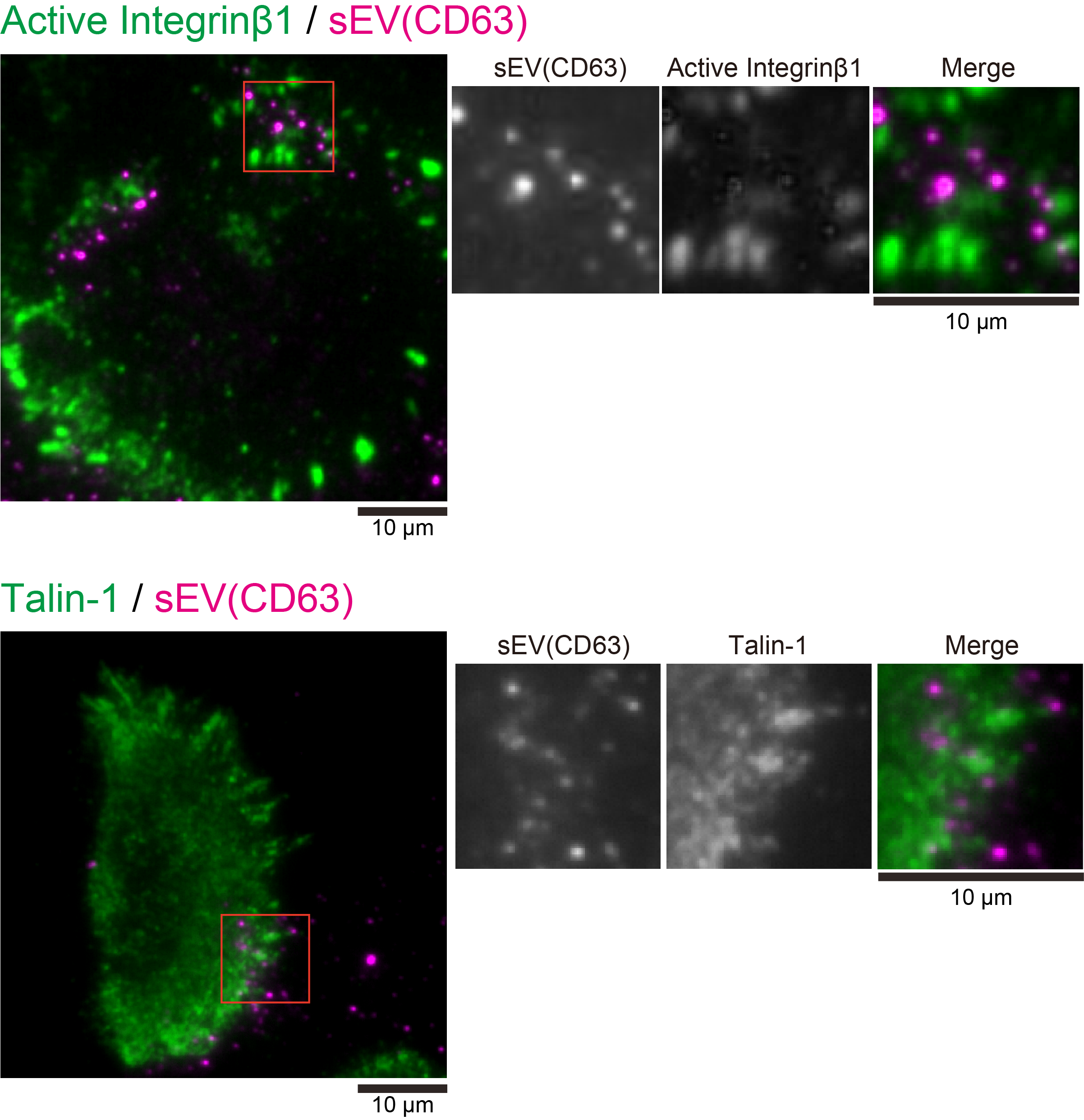
Supplementary Figure 12

**Supplementary Fig.12.** Immunostained images of sEVs containing CD63 and the active form of integrin (top) or talin-1 (bottom). Cells were incubated with sEVs containing CD63-Halo7 labeled with SF650T-Halo ligand (5.0 × 10^9^ particles/mL final concentration) for 1 h and then fixed with 4% paraformaldehyde. Subsequently, cells were incubated with antibodies against either the active form of integrin (HUTS-4) or talin-1 for 30 min, followed by staining with Alexa488-conjugated secondary antibody for mouse IgG. CD63-Halo7-SF650T in sEV particles was observed by TIRFM with single-molecule sensitivity. The active forms of integrin β1 and talin-1 labeled with Alexa488-conjugated secondary antibody were visualized by TIRFM at much lower sensitivity (488 nm laser intensity was 0.3 μW/μm^2^) to perform the ensemble-averaged imaging.

Supplementary Figure 13
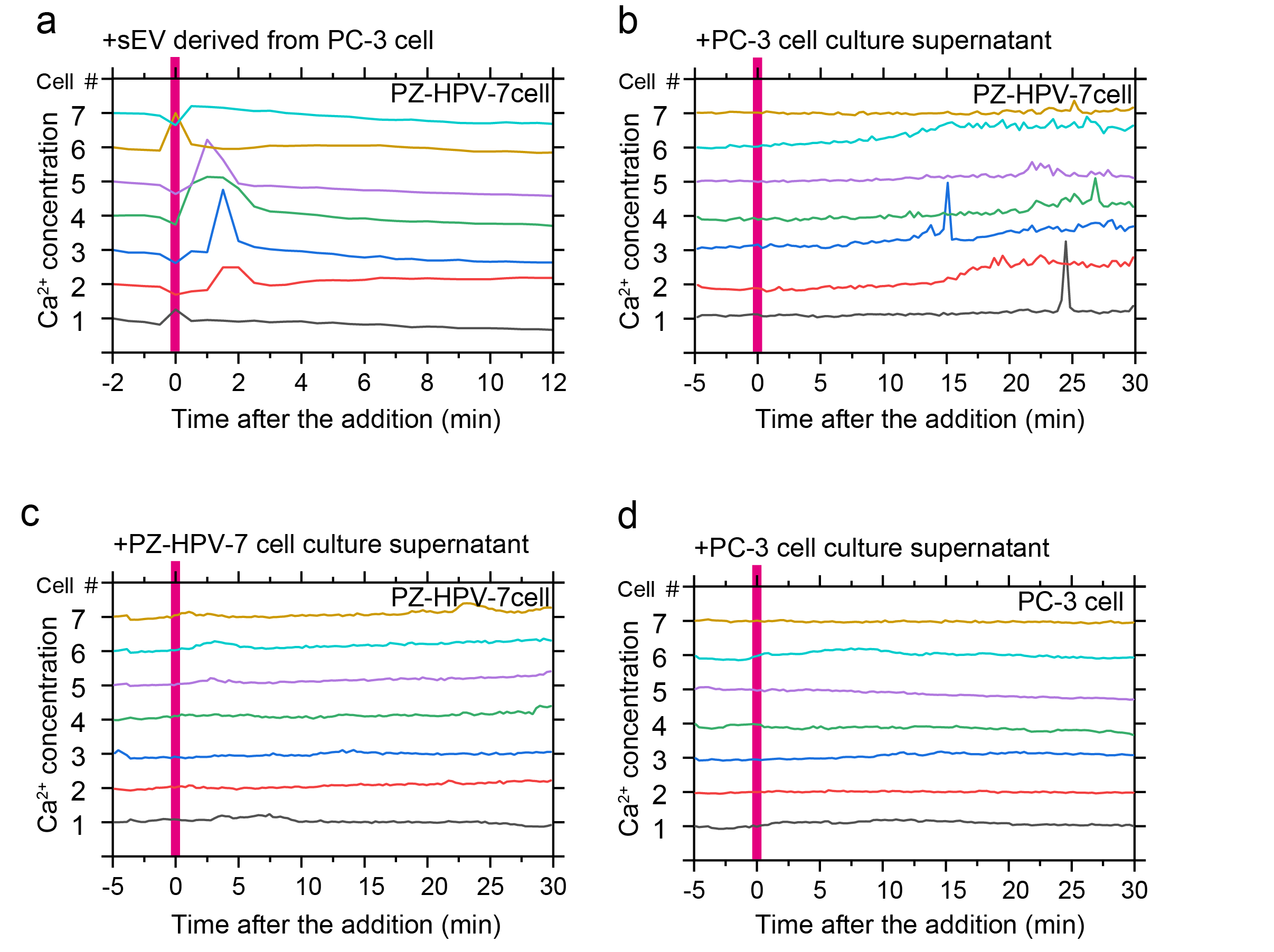

**Supplementary Fig.13.** Intracellular Ca^2+^ mobilization in response to the addition of sEVs or serum-free cell culture supernatant to recipient cells. Time courses of Fluo-8H fluorescence intensity, acquired by confocal microscopy, in PZ-HPV-7 cells after the addition of (**a**) sEVs derived from PC-3 cells, (**b**) unconcentrated serum-free PC-3 cell culture supernatant, and (**c**) unconcentrated serum-free PZ-HPV-7 cell culture supernatant. (**d**) Time course of Fluo-8H fluorescence intensity in PC-3 cells after the addition of unconcentrated serum-free PC-3 cell culture supernatant. Each panel presents data from seven representative cells.

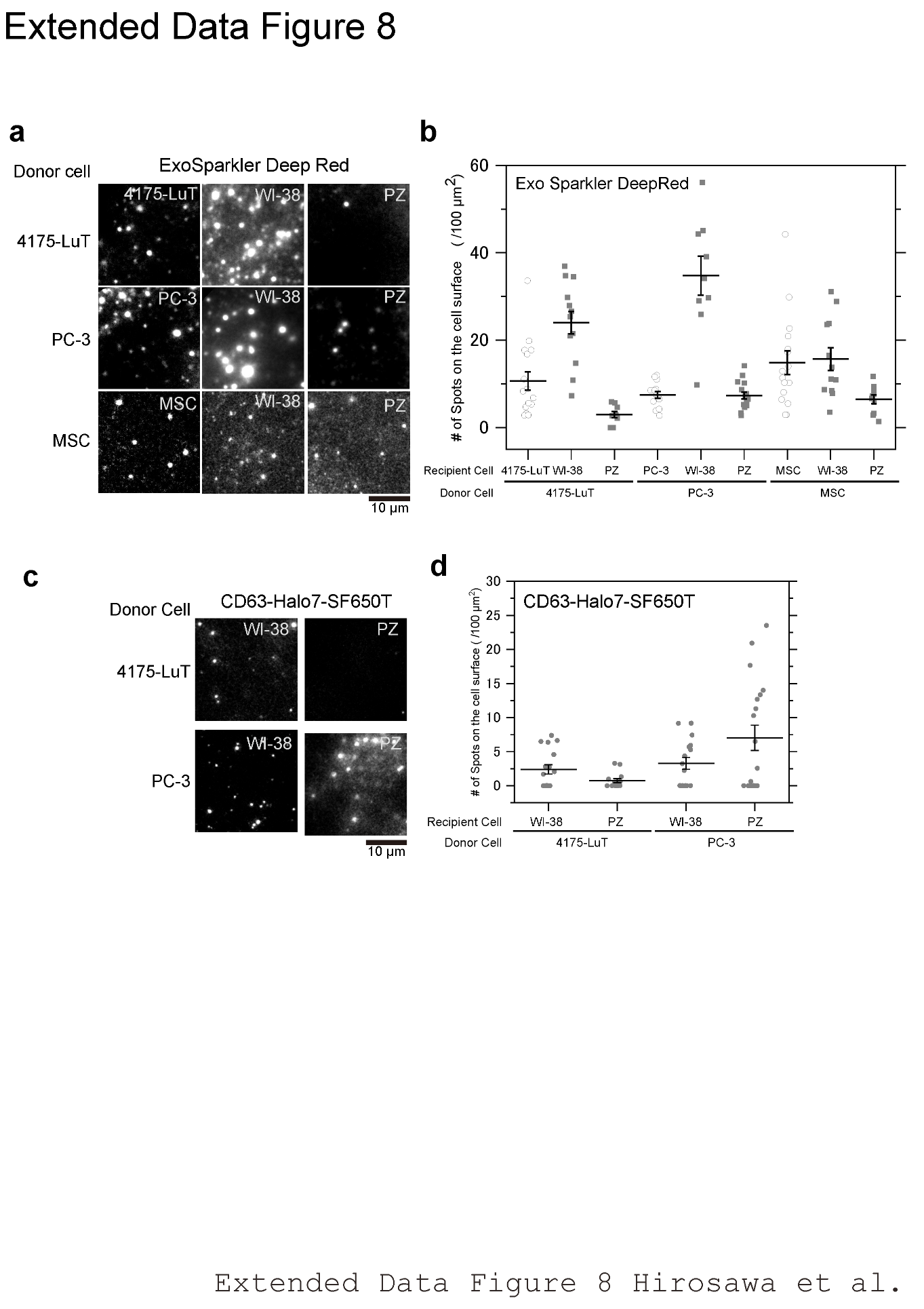
Supplementary Figure 14

**Supplementary Fig. 14.** The binding affinity of sEVs to recipient cells was dependent on the cellular origin of the sEVs and the recipient cells. **a**, Typical TIRF images of sEVs labeled with ExoSparkler Deep Red bound to recipient cells 30 min after sEV addition. **b**, Number of sEV particles bound to the recipient cell PMs. Autocrine combinations of sEVs and recipient cells are shown as open circles. Paracrine combinations are shown in closed rectangles. **c**, Typical TIRF images of sEV-CD63Halo7-SF650T attached to recipient cells 30 min after its addition. **d**, Number of sEV particles on the cell surface for each combination of sEVs and recipient cells.

Supplementary Figure 15

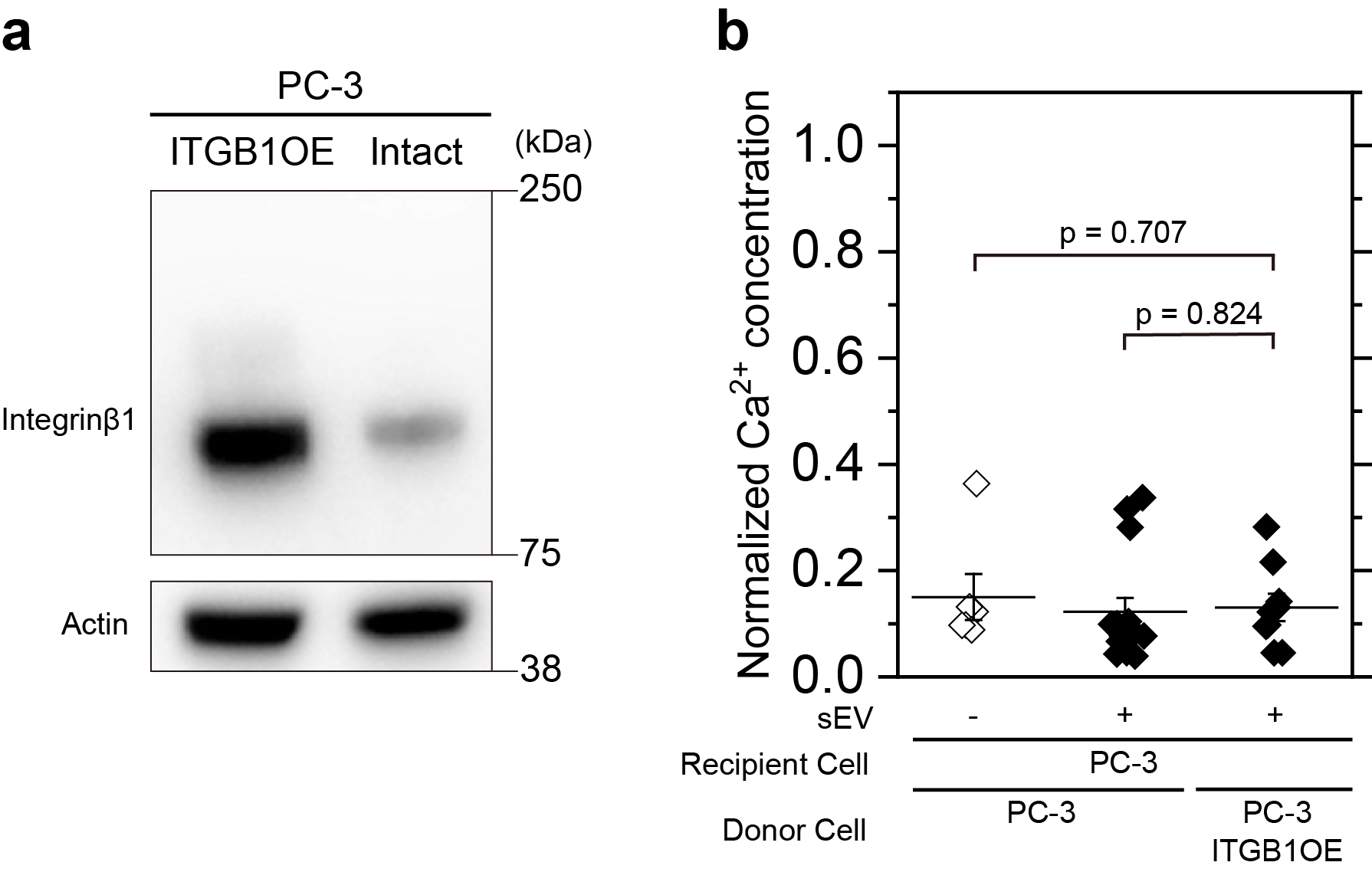

**Supplementary Fig. 15.** sEVs derived from donor PC-3 cells overexpressing integrin β1 (ITGB1OE) did not induce a Ca^2+^ response in the PC-3 recipient cells (autocrine). **a**, Western blot analysis showing integrin β1 overexpression in donor PC-3 cells. **b**, Normalized Ca^2+^ concentration after the addition of sEVs (5.0 × 10^9^ particles/mL final concentration) derived from intact or integrin β1-overexpressing donor PC-3 cells. Neither sEVs derived from intact nor integrin β1-overexpressing donor PC-3 cells induced an intracellular Ca^2+^ response in the recipient PC-3 cells.

Supplementary Figure 16

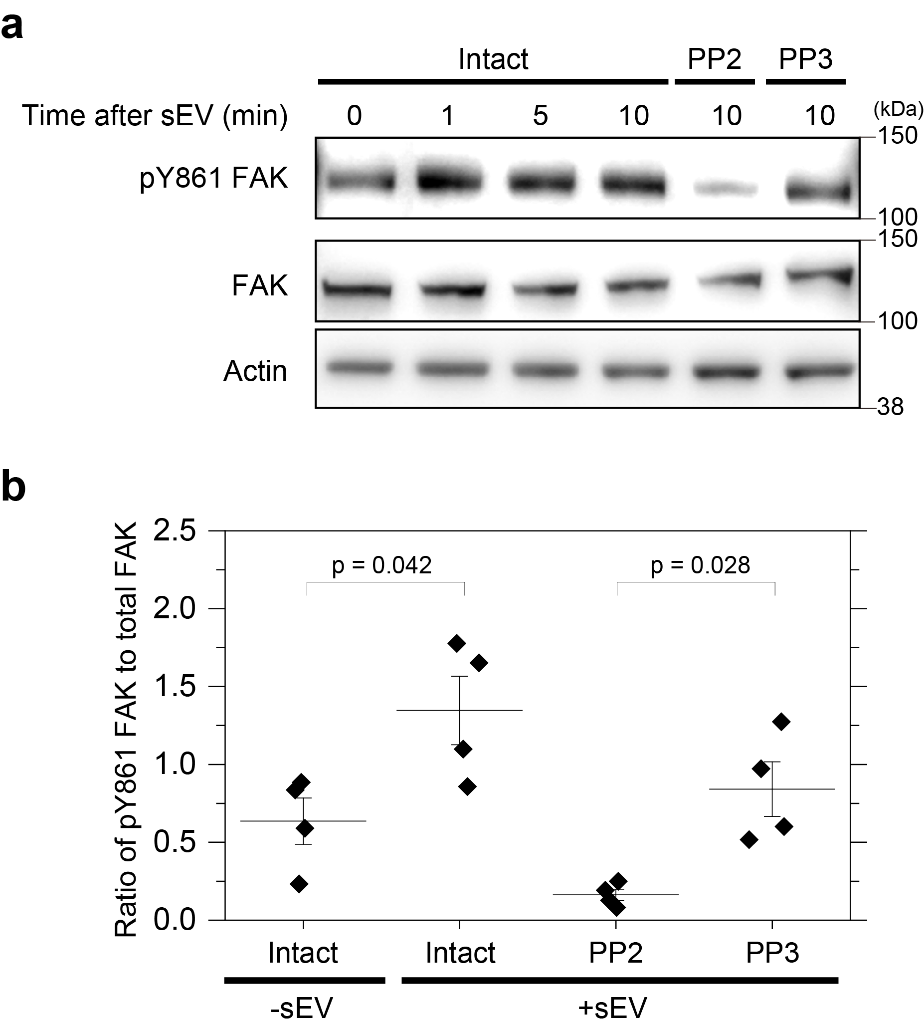

**Supplementary Fig. 16.** Focal adhesion kinase (FAK) phosphorylation after sEV addition to recipient PZ-HPV-7 cells and the effect of Src family kinase inhibition on phosphorylation. **a**, Western blot analysis showing changes in the phosphorylation level of tyrosine 861 (Y861) of FAK in recipient PZ-HPV-7 cells under various conditions: before and after the addition of sEVs derived from PC-3 cells and in the presence or absence of Src family kinase inhibitor (PP2) or its control analog (PP3). **b**, Ratio of phosphorylated Y861-FAK to total FAK (n = 4). These results indicate that sEVs derived from PC-3 cells induce Src family kinase-mediated FAK phosphorylation in recipient PZ-HPV-7 cells.

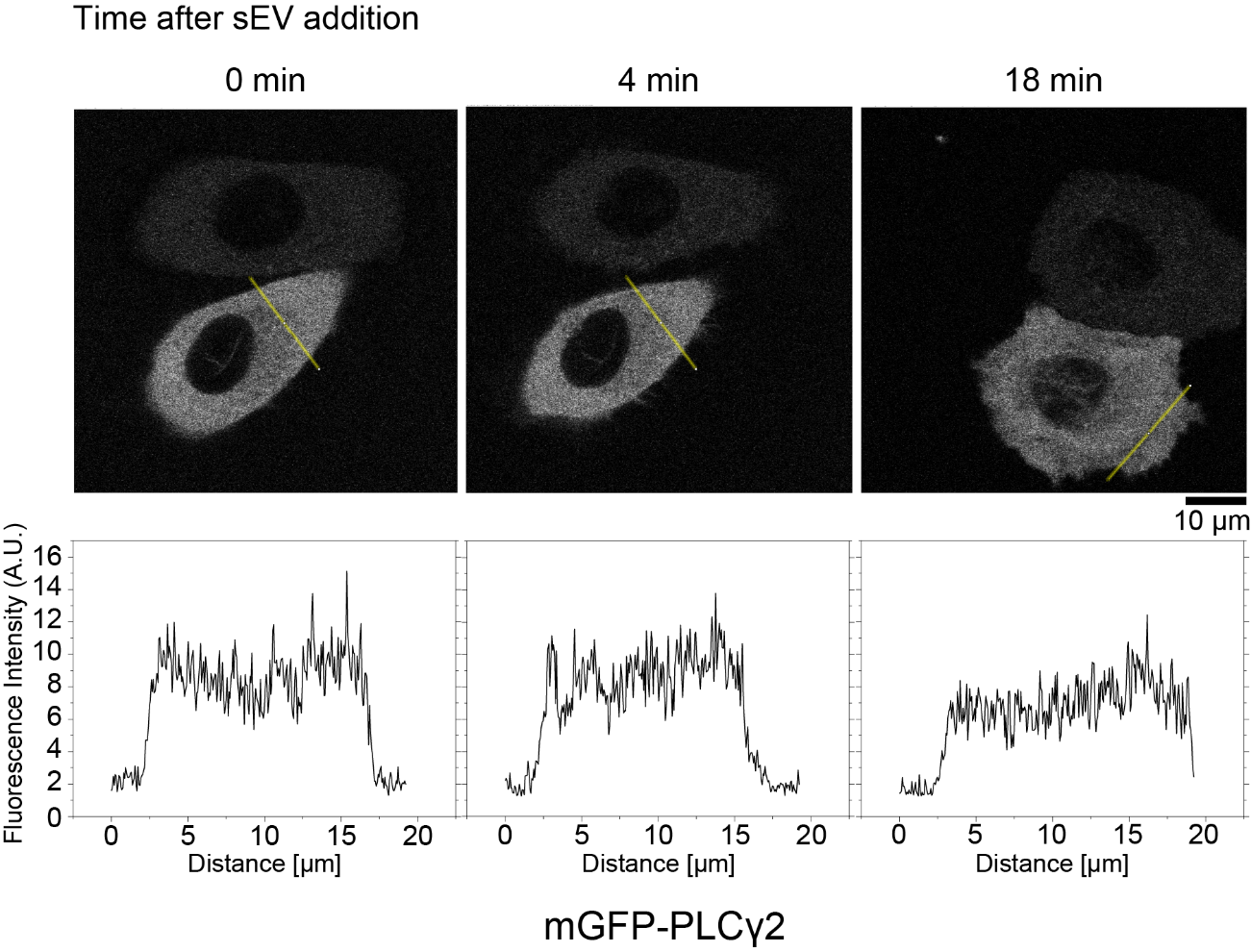
Supplementary Figure 17

**Supplementary Fig. 17.** Analysis of PLCγ2 distribution in recipient PZ-HPV-7 cells after the addition of PC-3 cell-derived sEVs by confocal microscopy. The distribution of mGFP-PLCγ2 fluorescence intensity in PZ-HPV-7 cells was measured before and after the addition of PC-3 cell-derived sEVs (top). Fluorescence intensity profiles along the yellow lines are shown at the bottom.

Supplementary Figure 18
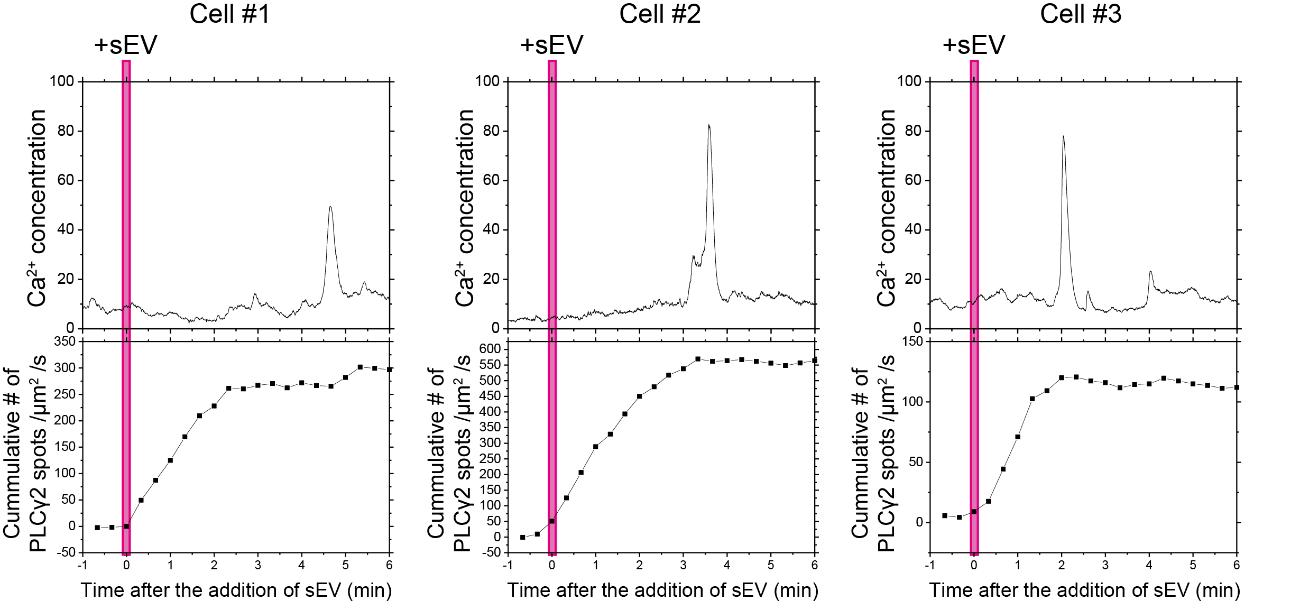

**Supplementary Fig. 18.** Temporal correlation between PLCγ2 recruitment to the PM and Ca^2+^ spikes after sEV addition. Time courses of Fluo-8H fluorescence intensity (top) and cumulative number of TMR-Halo7-PLCγ2 recruitment events (bottom) after sEV addition (5.0 × 10^9^ particles/mL final concentration) to the recipient cell. Fluo-8H was observed by oblique angle illumination, and single molecules of PLCγ2 were simultaneously monitored by TIRFM. The frequency of PLCγ2 recruitment was calculated as the deviation from the mean recruitment frequency before sEV addition. Examples from three individual cells are shown. In all cells, Ca^2+^ spikes occurred around the time when the cumulative number of PLCγ2 recruitment events to the PM reached a plateau. Combined with experiments with a PLC inhibitor (U73122), these results indicate that PLCγ2 recruitment triggers Ca^2+^ signaling after sEV addition.

Supplementary Figure 19

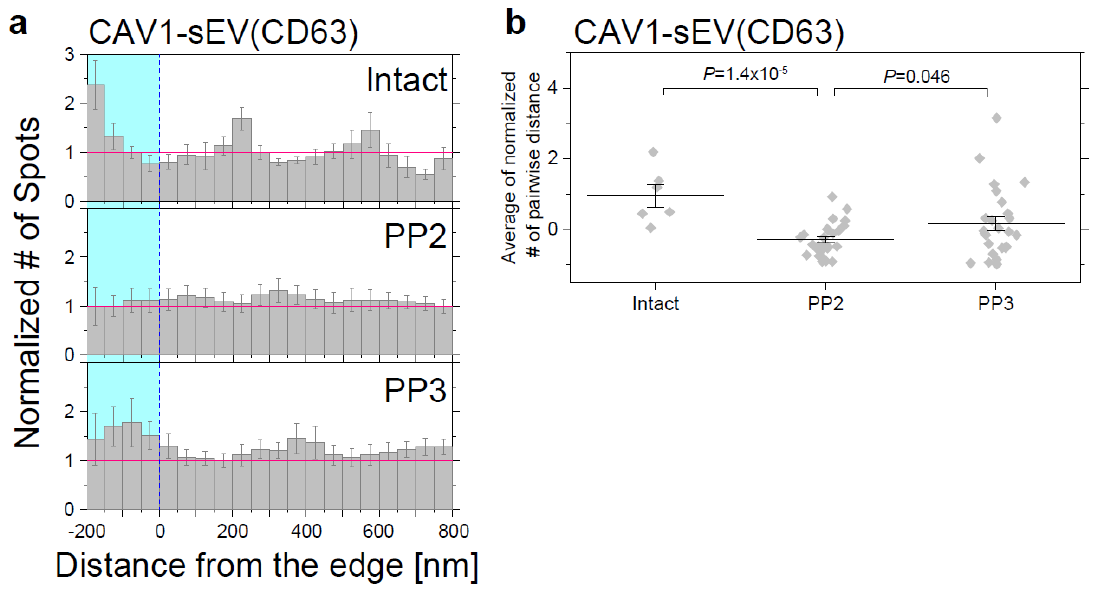

**Supplementary Fig. 19.** Src family kinase activity and caveolae-mediated sEV uptake. **a**, Probability density analysis of sEVs and caveolae in intact cells, cells treated with Src family kinase inhibitor (PP2), or its control analog (PP3). Simultaneous imaging of single sEV particles containing CD63-Halo7-SF650T and PALM movie observation of caveolae with CAV1-mEos4b was performed as in Fig. 4b. Colored areas indicate regions within caveolae. **b**, The average value of the normalized number of pairwise distances from -200 to 0 nm in Supplementary Fig. 19a was calculated for each cell and plotted in the graphs. Data are presented as means ± SE.

**Captions for supplementary movies**

**Supplementary Movie1** (the raw data for **Fig. 3b**)

(Left) The 3D trajectory of single sEV particle (sEV-CD63Halo7-SF650T derived from PC-3 cells) near the recipient cell (PZ-HPV-7 cell) PM. The color of the spot indicates the z-position of the sEV particle shown in the color bar. (Right) A source movie of the trajectory shown in the left was recorded at 30 Hz.

**Supplementary Movie 2** (the raw data for **Fig. 4d, e**)

A representative PALM movie of CAV1-mEos4b (green) was merged with a movie of single sEV particles (sEVs-CD63Halo7-SF650T) (magenta) on the living PZ-HPV-7 cell PM. Single molecules of Cav1-mEos4b and single particles of sEVs were observed at 200 Hz. The superimposed movie was created according to the method shown in Fig 4b and replayed at video rate (30 Hz).

**Supplementary Movie 3** (the raw data for **Fig. 4d, f**)

A representative PALM movie of LAMP2C-mEos4b (green) was merged with a movie of a single sEV particle (sEVs-CD63Halo7-SF650T) (magenta) on the living PZ-HPV-7 cell PM. Colocalization events are indicated by yellow arrowheads. Single molecules of LAMP2C-mEos4b and single particles of sEVs were observed at 200 Hz. The superimposed movie was created according to the method shown in Fig 4b and replayed at video rate (30 Hz).

**Supplementary Movie 4** (the raw data for **Fig. 6a upper**)

A typical movie, showing the recruitment of single molecules of TMR-Halo7-GPI (green) to a single particle of sEV-CD63Halo7-SaraFluor650T (magenta) on the living PZ-HPV-7 cell PM. Colocalization events were indicated by the yellow arrowhead. The movie was recorded at video rate and replayed in real-time.

**Supplementary Movie 5** (the raw data for **Fig. 6a bottom**)

A typical movie, showing single molecules of TMR-Halo7-Integrin β1 (green) and a single particle of sEV-CD81Halo7-SaraFluor650T (magenta) on the living PZ-HPV-7 cell PM. Colocalization events were indicated by the yellow arrowhead. The movie was recorded at video rate and replayed at 10 Hz

**Supplementary Movie 6** (the raw data for **Fig. 6c upper**)

A representative PALM movie of mEos4b-talin1 (green) was merged with a movie of a single particle of sEV-CD63Halo7-SaraFluor650T (magenta) on the living PZ-HPV-7 cell PM. Single molecules of mEos4b-talin1 and a single particle of CD63Halo7-SaraFluor650T were observed at 200 Hz. The superimposed movie was created according to the method shown in Fig. 4b and replayed at a video rate (30 Hz).

**Supplementary Movie 7** (the raw data for **Fig. 7a upper**)

The addition of PC-3-derived sEVs induced Ca^2+^ response in PZ-HPV-7 cells. The change in the intracellular Ca^2+^ concentration was monitored using the Fluo-8H indicator. sEVs were added to the cells at 2.5 minutes after the start of observation. Ionomycin was added to the cells at 20 minutes after the start of the observation. scale bar: 20 mm. The movie was recorded at 0.05 Hz and replayed at 30 Hz.

**Supplementary Table 1.** cDNA used in this study

| Protein name | Species | Source | Tag | Tag position | Linker sequence | Vector |
| --- | --- | --- | --- | --- | --- | --- |
| CD9 | Human | Kazusa DNA research institute (FHC08599) | mEGFP | C terminus | TGGGRASGGGSGGSGGGSGGSGGGSGG | pcDNA6 |
| CD9 | Human | Kazusa DNA research institute (FHC08599) | Halo7 | C terminus | TGGGRASGGGSGG | pEGFPN1 |
| CD9 | Human | Kazusa DNA research institute (FHC08599) | Halo7 | C terminus | TGGGRASGGGSGG | pPBZeo |
| CD63 | Human | Kazusa DNA research institute (FHC02909) | mEGFP | C terminus | TGGGRASGGGSGGSGGGSGGSGGGSGG | pcDNA6 |
| CD63 | Human | Kazusa DNA research institute (FHC02909) | Halo7 | C terminus | TGGGRASGGGSGG | pEGFPN1 |
| CD81 | Human | Kazusa DNA research institute (FHC08124) | mEGFP | C terminus | TGGGRASGGGSGGSGGGSGGSGGGSGG | pcDNA6 |
| CD81 | Human | Kazusa DNA research institute (FHC08124) | Halo7 | C terminus | TGGGRASGGGSGG | pEGFPN1 |
| CD81 | Human | Kazusa DNA research institute (FHC08124) | Halo7 | C terminus | TGGGRASGGGSGG | pPBZeo |
| AP2alpha1v2 | Human | Kazusa DNA research institute (FHC03782) | mEos4b | C terminus | PRARDPPVAT | pEGFPN1 |
| CAV1 | Human | Kazusa DNA research institute (FHC04442) | mEos4b | C terminus | PRARDPPVAT | pEGFPN1 |
| LAMP2C | Human | Kazusa DNA research institute (FHC07596) | mEos4b | C terminus | PRARDPPVAT | pEGFPN1 |
| LAMP2C | Human | Kazusa DNA research institute (FHC07596) | Halo7 | C terminus | SGGGGSGGGGSGGGG | pEGFPN1 |
| Dynamin(K44A) | Rat | Addgene (#22301) | EGFP | C terminus | AAGDSVLGGILQSTVPRARDPPVAT | pEGFPN1 |
| GPI　(CD59) | Human | Previous paper in our group | Halo7 | N terminus | N/A | pER |
| GPI　(CD59) | Human | Previous paper in our group | mGFP | N terminus | N/A | pER |
| TM　(LDLR) | Human | Previous paper in our group | Halo2 | N terminus | N/A | pEGFPN1 |
| Integrin beta1 | Human | Slightly modified with Previous paper ^7^) | Halo7 | Extracellular loop of integrin (insert between Gly101 and Tyr102) | EFGGSGGSG /GGSGGSGLE | pEGFPN1 |
| Integrin beta1 | Human | Previous paper ^7^ | - | - | - | pPBpuro |
| PLC gamma 1 | Rat | a gift from Dr. Matilda Katan | EGFP | C terminus | ADPPVAT | pEGFPC1 |
| PLC gamma 2 | Human | a gift from Dr. Matilda Katan^8^ | mEGFP | N terminus | SGLRSRAQASNSRPT | pEGFPC1 |
| PLC gamma 2 | Human | a gift from Dr. Matilda Katan^8^ | Halo7 | N terminus | SGLRSRAQASNSRPT | pEGFPC1 |
| Talin-1 | Human | a gift from Dr. Akihiro Kusumi^9^ | mEos4b | N terminus | SGLRSRAQASSGGGGSGGGGSGGGGTSQACRRIL | pEGFPN1 |
| Talin head | Human | This paper (Modified from Talin-1) | mEGFP | N terminus | SGLRSRAQASSGGGGSGGGGSGGGGTSQACRRIL | pEGFPN1 |
| FAK | Mouse | a gift from Dr. Caroline H. Damsky^10^ | mEos4b | N terminus | SGGGGSGGGGSGGGG | pEGFPN1 |
| Rab11 | Human | a gift from Dr. Kazuhisa Nakayama^11^ | mEos4b | N terminus | SGGGGSGGGGSGGGG | pEGFPN1 |
| piggyBacTransposase | - | Previous paper ^12, 13^ | - | - | - | pCMV-mPBase |
| Rab6a | Mouse | a gift from Dr. Tomohiko Taguchi^14^ | mEos4b | N terminus | GGSGGGSGGGS | pMX-IRES |
| STING | Mouse | a gift from Dr. Tomohiko Taguchi^14^ | mEos4b | N terminus | GGGGSGGGGSGGGGS | pMX-IRES |
